# Mutation Profiles, Glycosylation Site Distribution and Codon Usage Bias of HPV16

**DOI:** 10.1101/2021.03.04.434005

**Authors:** Wei Liu, Junhua Li, Hongli Du, Zhihua Ou

## Abstract

Human papillomavirus type 16 (HPV16) is the most prevalent HPV type causing cervical cancers. Herein, using 1,597 full genomes of HPV16, we systemically investigated the mutation profiles, surface protein glycosylation sites and the codon usage bias of the eight open reading frames (ORFs) of HPV16 genomes from different lineages and sublineages. Multiple lineage- or sublineage-specific mutation sites were identified. Glycosylation analysis showed that HPV16 lineage D contained the highest number of unique potential glycosylation site in both L1 and L2 capsid protein, which might lead to their antigenic distances from other HPV16 lineages. Nucleotide composition of HPV16 showed that the overall AT content was higher than GC content at the 3^rd^ codon position. Relatively high ENC values suggested that the HPV16 ORFs didn’t have strong codon usage bias. Most of the HPV16 ORFs were mainly governed by natural selection pressure such as translational pressure, except for L2. HPV16 only shared some of the preferred codons with human, which might help reduce competition in translational resources. These findings may help increase our understanding of the heterogeneity between HPV16 lineages and sublineages, and the adaptation mechanism of HPV in human cells, which might facilitate HPV classification and improve vaccine development and application.

## 1. Introduction

Human papillomaviruses (HPVs) can cause mucous and cutaneous infections. Up to now, more than 200 different HPV types have been identified (http://www.hpvcenter.se/). According to their carcinogenicity, HPVs can be divided into high-risk and low-risk types. High-risk types include HPV16, 18, 31, 33, 35, 39, 45, 51, 52, 58, 59, etc. [1], which mainly cause reproductive tract diseases. Among them, HPV16 is the dominant type leading to cervical cancer and accounts for above 50% cervical cancer cases [2, 3].

HPVs are double-stranded circular DNA viruses with a genome size of about 8kb. HPV16 genomes include three general regions: a region encoding early-stage proteins (E1, E2, E4, E5, E6 and E7), a region encoding late-stage proteins including L1 and L2, and an upstream regulatory region (URR) [4]. E1 and E2 proteins regulate the replication and transcription of HPV genomes [5, 6]. E5, E6 and E7 proteins are cofactors for HPV carcinogenesis, involving in epithelial dysplasia and tumor progression after HPV infection [7–10]. L1 and L2 are the major and minor capsid proteins, which are expressed during the late stage of HPV infection. Besides forming the elegant icosahedral surface of the papillomavirus virion, these two capsid proteins are essential for virus binding and entry into cells [11, 12]. Currently, L1 and L2 proteins are the targets of HPV prophylactic vaccines, while E6 and E7 are targets of therapeutic vaccines of HPV-induced cancers [13].

Above the type or genotype level, HPVs are classified based on the nucleotide sequence of L1 [14, 15]. In 2013, Chen et al. proposed the lineage/sublineage classification criteria for papillomaviruses of the same type based on the nucleotide difference of complete genomes, with 1.0-10.0% and 0.5-1.0% differences defining different lineages and sublineages [16]. Up to date, four lineages (A-D) and sixteen sublineages (A1−4, B1−4, C1−4 and D1−4) have been identified for HPV16 around the world [17, 18].

It has been reported that HPV16 sublineages differ in their geographic distribution and carcinogenicity [19–21]. A1 sublineage was the dominant sublineage in Europe, the Americas, South Asia and Oceania, and A2 sublineage was distributed in Europe, North America and Oceania, while A3 and A4 sublineages were mainly distributed in East Asia. Lineage B and C were almost exclusively distributed in Africa, and lineage D was the most common in South/Central America [22]. Mirabello et al. found that white women infected with HPV16 A1/A2 variants had an increased risk of CIN3+ progression, and A4 sublineage was associated with an increased risk of cancer in Asian women [23].

Glycosylation plays an important role in the folding and stabilization of viral proteins, recognition of host cell receptors and immune escape of viruses. Mutation of the N-glycosylation site of the surface envelope glycoprotein of HIV, gp120, would remove the glycosylated oligosaccharide chain and expose the masked antigenic epitopes, increasing the antigenic recognition of gp120 by the antibodies [24]. Addition of glycosylation to the hemagglutinin and neuraminidase protein of influenza viruses can result in viral antigenic drift from older strains [25, 26]. Therefore, it is meaningful to elucidate the potential glycosylation modification sites of viral surface proteins, which could help uncover novel molecular determinants of antigenic differences and improve vaccine design.

A trinucleotide codon is used to encode one standard amino acid, and most amino acids are coded by more one codon, except Met and Trp. The codons code for the same amino acid are called synonymous codons. Usage of synonymous codons may vary between and within species, which is called codon usage bias (CUB). Natural selection, mutation pressure, and other factors can all affect CUB [27, 28]. Viruses depend on their host for survival, so their codon usage patterns may be similar to those of the host in order to express viral proteins efficiently [27, 29]. However, it has also been found that some viruses may have CUB different from their host to escape from the host immune system [30]. It has been shown that the genera *Alphapapillomavirus* and *Betapapillomavirus* had different CUB, and the different codon usage pattern may be related to the histological specificity of the papillomaviruses [31]. CUB was correlated with high A + T content at the 3^rd^ codon position of HPV genes [32]. Codon-optimized HPV16 E6 and E7 genes were suggested for the development of therapeutic vaccines against HPV16 [33, 34] to improve protein production level in cells. Understanding the CUB of HPV16 genes would also increase our understanding in their interaction with human hosts and the mechanism underlining persistent infection.

The rapid accumulation of HPV16 genome data has provided a new opportunity for extensive and in-depth research on the genetic diversity of HPV16. In this study, we aimed to explore the genomic mutation profiles and the glycosylation site distribution for surface proteins in different HPV16 sublineages. The subsequent findings would help us further understand the heterogeneity between the sublineages and how such differences might influence surveillance and vaccine application. To further understand the viral-host interaction mechanism of HPV16, we also comprehensively analyzed the codon usage patterns of the eight HPV16 ORFs and compared their viral CUB with that of humans.

## 2. Materials and Methods

### 2.1 Data preparation

A total of 3,729 complete sequences of HPV16 genomes were retrieved from the National Center for Biotechnology Information (NCBI) (http://www.ncbi.nlm.nih.gov/Gen-bank/) as of May 13, 2020. In order to get high quality genomes, these sequences were processed as follows: (1) sequences with a length of 7000-8500bp and ambiguous sites less than 5 were kept; (2) sequences that contain 70 or more consecutive “N” (about 1% complete sequence) were removed; (3) sequences were aligned by MAFFT v7.407 [35]; (4) the aligned sequences were checked in BioEdit v7.0.5 [36] and low-quality sequences and those with early stop codons were removed. DNA sequences were translated into amino acid sequences to ensure correct reading frames. Finally, a total of 1,597 genomes were included for this study. The eight ORFs were extracted based on the coordinate of HPV16 reference genome (Accession Number K02718), considering that the starting position of most E6 genes is 104, it was set as the starting position of all E6 genes. The detailed information of the genomes, such as host origins, geographical locations and collection time, were provided in Table S1.

### 2.2 Phylogenetic and cluster analysis

In order to compare the codon usage preferences between different sublineages, the whole genomes obtained were classified into different sublineages. Maximum likelihood phylogeny was constructed with IQ-TREE using TVM+F+I+G4 nucleotide substitution model with 1000 ultrafast bootstrap implementation [37–39]. The nucleotide difference between all sequences and the reference sequences were calculated with R package seqinr v3.6-1. According to the phylogenetic topology and sequence differences (inter-lineage difference: 1%-10%; inter-sublineage difference: 0.5%-1%), all sequences were divided to lineages and sublineages for downstream analysis. The reference sequences of different lineages/sublineages were obtained from GenBank [18], with their accession numbers as follows: K02718 (A1), AF536179 (A2), HQ644236 (A3), AF534061 (A4), AF536180 (B1), HQ644298 (B2), AF472509 (C1), HQ644257 (D1), AY686579 (D2), AF402678 (D3), HQ644244 (C2), KU053920 (C3), KU053925 (C4), KU053931 (D4), KU053915 (B3), KU053914 (B4).

### 2.3 Mutation detection of ORFs

Nucleotide sequences of the eight ORFs were compared against the reference genome (K02718) to identify mutations. The amino acid mutations resulting from the nucleotide mutation were also determined.

### 2.4 Identification of potential glycosylation sites in L1 and L2 protein

L1 and L2 sequences were translated into protein sequences in BioEdit. The potential glycosylation sites were determined by identification of N-linked glycosylation motifs (N-X-T/S, X: any amino acid except for P) in the protein sequences.

### 2.5 Nucleotide composition analysis

Calculations of the GC content at the 1^st^, 2^nd^ and 3^rd^ codon positions (GC1, GC2, GC3) and the average content of GC1, GC2 (GC12) of all ORFs were conducted with R package SADEG v1.0.0 [40].

### 2.6 Analysis of effective number of codons

Effective number of codons (ENC) is a parameter to evaluate the overall codon preference of genes. ENC value ranges from 20 to 61. The value of 20 means that only one codon is used for each amino acid, and 61 means that every codon is used [41]. The lower the ENC value, the stronger the bias for codon usage. Genes with low expression levels were found to have high ENC values and more rare codons [42]. The ENC was calculated using R package SADEG. The formula for calculating ENC value is as follows [41]:

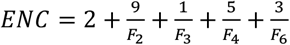

where F2 is the identical probability of two synonymous codons randomly selected.

Wright [41] suggested that the ENC-plot (ENC plotted against GC3) could be used to investigate codon usage patterns across genes, and that ENC value is independent of gene length and amino acid (aa) composition. The standard curve in ENC-plot analysis represents that CUB is completely determined by nucleic acid composition. If a point falls on the expected curve, the codon usage is influenced by mutational pressure; If a point falls below the standard curve, its codon usage is also affected by selection pressure. The expected ENC was calculated using the equation:

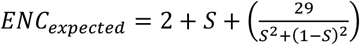

where S indicates the content of GC3.

### 2.7 Neutrality plot analysis

Both mutational pressure and natural selection affects the bias of codon usage. Amino acid changes at the 3^rd^ codon positions usually cause synonymous mutation, which indicate a mutational pressure, while nucleotide changes causing nonsynonymous mutations indicate selection pressure. The ratio of GC12 and GC3 is used to measure the influence of natural selection and mutation pressure. The slope of the regression line represents the evolutionary speed of the mutation pressure and natural selection pressure [43]. The more the slope close the diagonal of the coordinate axis, the greater the influence of the mutation pressure. However, if the regression line deviates from the diagonal, the selected codons were influenced by other factors, like natural selection [44].

### 2.8 Codon usage frequency analyses

Relative synonymous codon usage (RSCU) is largely independent of amino acid composition and can be used to compare codon usage among genes or genomes with different lengths and amino acid compositions. The calculation of the RSCU value assumes that the codons of the same specific amino acid have equal usage, and the ratio of the actual codon usage frequency to the expected frequency is defined as the RSCU value [45]. RSCU values of <0.6, 0.6-1.6, >1.6 indicate low, normal, over usage of the codon [44]. The average RSCU data of human was originated from work by Malik et al. [43], while the mean RSCU values of HPV16 ORFs were calculated by R package SADEG v1.0.0. [40]. The calculation formula of RSCU is as follows,

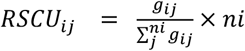

where i is i-th codon and j is j-th amino acid, gij is the observed number of the i-th codon for the j-th amino acid that has an “ni” type of synonymous codon [45].

## 3. Results

### 3.1 Classification of HPV16 lineages and sublineages

Using 1,597 full genomes (Supplementary Table S1), we constructed a Maximum Likelihood tree (Supplementary Figure S1) and conducted lineage/sublineage classification based on the criteria proposed by Chen et al [16]. Only one sequence was not assigned to any lineage/sublineage because its long distance to other known lineages. In summary, we obtained 1352 (84.7%) sequences from lineage A, 34 (2.1) from lineage B, 56 (3.5%) from lineage C, and 154 (9.6%) from lineage D (Supplementary Table S2). Of all the sequences in lineage A, 1,053 (77.9%) genomes belonged to A1 sublineage (Table 1, Supplementary Table S2), following by A2 (204), A4 (84) and A3 (11) sublineages. Unfortunately, the number of genomes in several B and C sublineages were less than 5 sequences. Other sublineages with more than 10 sequences included B1 (28), C1 (50), D1 (12), D2 (35), D3 (95) and D4 (12).

**Table 1.** Mutation profiles of HPV16 sublineages.

| ORF | Nucleotide<br>mutation | Amino acid<br>mutation | Proportion of sequences with the corresponding mutations in each sublineage (%) |  |  |  |  |  |  |  |  |  |
| --- | --- | --- | --- | --- | --- | --- | --- | --- | --- | --- | --- | --- |
|  |  |  | A1 | A2 | A3 | A4 | B1 | C1 | D1 | D2 | D3 | D4 |
|  |  |  | (n=1053) | (n=204) | (n=11) | (n=84) | (n=28) (B,<br>n=34) | (n=50) (C,<br>n=56) | (n=12) | (n=35) | (n=95) | (n=12) |
| **E1** | A1931C | E399D |  |  | **90.9** |  |  |  |  |  |  |  |
|  | G2336A | M534I | 1.9 |  | 9.1 | **97.6** |  |  |  |  |  |  |
|  | G2649A | E639K | 1 |  |  |  |  |  |  | **100** | **98.9** |  |
| **E2** | A3180C | E142D |  |  |  |  | 14.3 (14.7) | 6 (7.1) |  |  | **97.9** |  |
|  | C3158G | T135R |  |  |  |  |  |  |  |  |  | **100** |
|  | G3412A | A220T |  |  |  |  |  |  | **100** |  |  |  |
|  | G3415A | A221T |  |  |  |  |  |  |  | **100** |  |  |
|  | G3430A | A226T |  |  |  |  |  | **100 (98.2)** |  | 2.9 |  |  |
|  | T3223A | L157I/M<sup>a</sup> |  |  |  |  |  |  | **100** | **100** | **100** | **100** |
|  | T3383C | I210T | 1.7 | **100** |  | **100** |  |  |  |  |  |  |
|  | T3386C | I211T |  |  |  |  |  |  |  |  | **95.8** |  |
| **E5** | A4054T | I65L |  |  |  |  | **100 (100)** |  |  |  |  |  |
|  | G3881A | A7T |  |  |  |  | **100 (100)** |  |  |  |  |  |
| **E6** | G132T | R10I |  |  |  |  |  | **98 (87.5)** |  |  |  |  |
|  | T350G | L83V | 47.8 | 21.6 |  |  | 3.6 (14.7) |  | **100** | **100** | **100** | **100** |
| **E7** | A647G | N29S |  |  |  | **98.8** |  | **100 (89.3)** |  |  |  |  |
| **L1** | A6178C | N207T |  |  |  | 41.7 | 14.3 (11.8) | 78 (75) | 8.3 | 5.7 | 100 |  |
|  | A6801T | T415S |  |  |  |  |  |  |  |  | **97.9** |  |
|  | T6480C | S308P |  |  |  |  | 3.6 (2.9) | **100 (100)** |  |  |  |  |
| **L2** | A4967G | T245A |  |  |  |  |  |  | **100** | **97.1** | **98.9** | **100** |
|  | A5032T | L266F |  |  |  |  |  |  | **100** | **100** | **100** | **100** |
|  | A5288C | T353P |  |  |  |  |  | **100 (89.3)** |  |  |  |  |
|  | A5288G | T353A |  |  |  |  | **100 (97.1)** |  |  |  |  |  |
|  | T5366G | S379V/A<sup>b</sup> |  |  |  |  |  |  | **100** | **97.1** | **96.8** | **100** |
|  | T5384G | S385A |  |  |  |  |  |  | **100** | **97.1** | **100** | **100** |

**Note:** mutation sites were determined for sublineages with more than 10 sequences, and only those mutations occurred in >90% of the sequences in a certain sublineage were showed. Blank space indicates that there were little/no corresponding mutations in the sublineage or that sublineage contained less than 10 sequences. Because multiple sublineages of B and C lineage contained less than 10 strains, therefore, mutation frequencies were also calculated for B and C lineage. The numbers in parentheses indicate the proportion of the mutation in B or C lineage. Mutation frequencies over 85% are in bold.
<sup>a</sup> L157I/M: T3223A -> L157I; T3223A and A3224G -> L157M.
<sup>b</sup> S379V/A: T5366G -> S379A; T5366G and C5367T -> S379V.

### 3.2 Mutations identified across the HPV16 genome

Because different HPV sublineages displayed heterogeneity in geographical distribution and carcinogenic ability, we sought to identify mutations that significantly differ between the lineages and sublineages. Sites in the ORFs that differ from the reference sequence (K02718) were identified as mutation sites. The distributions of mutations by gene are shown in Figure S2. The L2 and E2 ORFs of HPV16 showed higher levels of genomic diversity than other genes, with 6,459 and 6,320 mutations detected in E2 and L2, respectively, while E7 was relatively conserved, with only 183 mutations observed (Supplementary Table S3, Figure S2). To identify lineage-specific genetic changes, mutations occurring in over 90% sequences of the sublineages that contained more than 10 sequences were further identified. There were at least 25 nucleotide sites displayed lineage-fixation in at least one sublineage (Table 1; Supplementary Table S3). Mutations including E2 T3223A, L2 A4967G, L2 A5032T, L2 T5366G and L2 T5384G were uniquely associated with lineage D, while E5 A4054T, E5 G3881A, L2 A5288G were uniquely associated with lineage B or sublineage B1, and E6 G131T and L2 A5288C were associated with sublineage C or sub-lineage C1. Several other mutations were found to be sublineage specific, including E1 A1931C for A3, E2 G3412A for D1, E2 G3415A for D2, E2 T3386C and L1 A6801T for D3, and E2 C3158G for D4. These mutations may be useful for the lineage or sublineage identification based on nucleotide polymorphism.

### 3.3 Glycosylation analysis of HPV16 L1 and L2 proteins

The L1 protein plays a major role in the receptor binding of HPVs to host cells [5]. L1 protein is the main component of the current HPV prophylactic vaccines, and the variation of its protein sequence is closely related to the effectiveness of the vaccines [46]. Due to the complex design of the multivalent L1-VLP vaccines, the vaccines cannot prevent all types of HPV infection, and some HPVs that can cause mucosal cancer cannot be covered. While the minor capsid protein L2 contains common epitopes that induce low titers of antibody, it can produce broadly cross-neutralizing antibodies against heterologous HPV types and might be served as a potential common HPV vaccine antigen [47]. To explore the variations of HPV16 L1 and L2 proteins, the amino acid sequences of L1 and L2 of 1,597 HPV16 genomes were predicted for glycosylation sites. The A1 sublineage had the largest number of potential glycosylation sites in L1 and L2 protein, which may be due to the abundant sequences within this sublineage (Supplementary Table S4 and Table S5). Ten and twenty-nine glycosylation sites were identified in all lineages for L1 and L2 proteins, respectively (Figure 1). Some glycosylation sites were lineage specific. In L1 protein, 27 glycosylation sites were observed only in A lineage, 1 in C lineage and 10 in D lineage. In L2 protein, 61 glycosylation sites were only found in A lineage, 2 in B lineage and 11 in D lineage. Collectively, the L1 and L2 glycosylation sites in lineage D displayed the largest differences from the other lineages, especially lineage A. These lineage-specific glycosylation sites may play an important role in host cell recognition and immune escape process.

**Figure 1.**
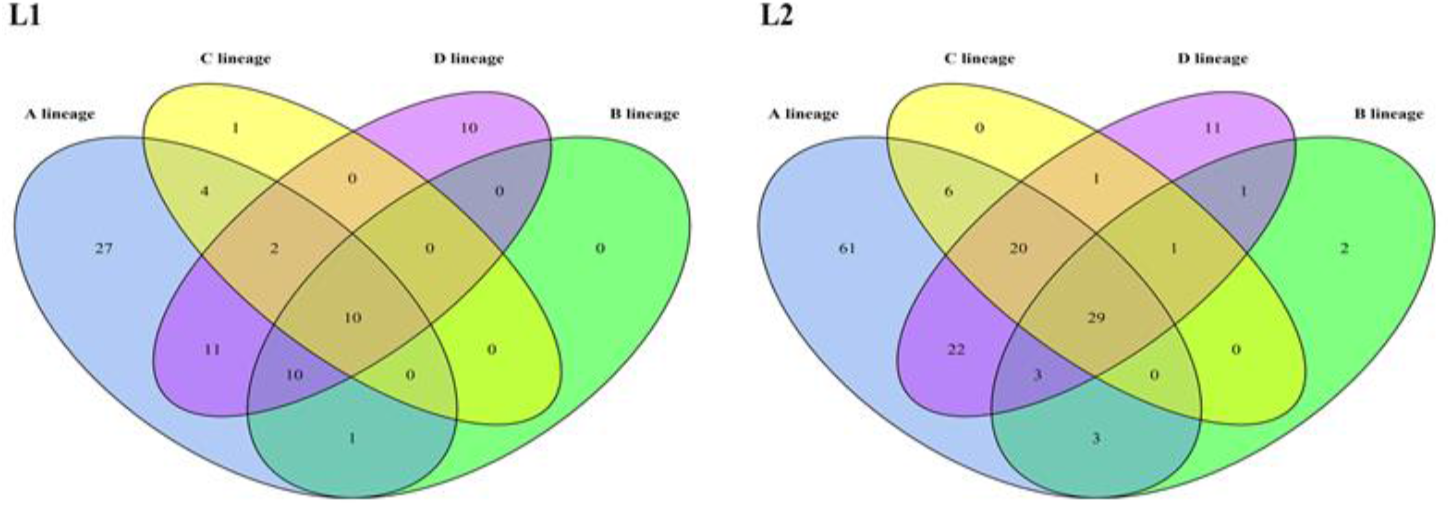
The lineage distribution of potential glycosylation sites on L1 and L2 proteins.

### 3.4 Nucleotide composition of HPV16 genomes

Our analysis on nucleotide contents showed that HPV16 genomes are AT-rich (Table 2). The mean nucleotide content of A and T for the eight ORFs (E1, E2, E4, E5, E6, E7, L1, L2) were 31.91% and 28.84%, respectively, higher than that of C and G. The mean G+C% of the eight ORFs ranged from 33.46% (E5) to 50.11% (E4). Comparison by codon positions showed that the third codon positions contained low GC content (15.07%-41.85%), with E1 (18.62%) and L2 (15.07%) showing extremely low values. These indicated that the third codon position mainly accounted for the nucleotide composition bias of HPV16.

**Table 2.** Nucleotide composition of the eight ORFs of HPV16 (%).

|  | T | C | A | G | A+T | G+C | GC1 | GC2 | GC12 | GC3 |
| --- | --- | --- | --- | --- | --- | --- | --- | --- | --- | --- |
| E1 | 28.27 | 12.88 | 36.40 | 22.45 | 64.67 | 35.33 | 41.96 | 36.83 | 39.40 | 18.62 |
| E2 | 26.00 | 18.38 | 36.34 | 19.27 | 62.35 | 37.65 | 46.24 | 37.22 | 41.73 | 27.88 |
| E4 | 18.94 | 31.43 | 30.93 | 18.70 | 49.87 | 50.13 | 54.43 | 52.42 | 53.43 | 41.85 |
| E5 | 43.37 | 17.90 | 23.16 | 15.56 | 66.54 | 33.46 | 32.90 | 35.71 | 34.30 | 24.49 |
| E6 | 28.17 | 15.65 | 34.18 | 22.00 | 62.35 | 37.65 | 42.07 | 39.16 | 40.62 | 31.21 |
| E7 | 25.36 | 20.85 | 30.92 | 22.87 | 56.28 | 43.72 | 54.73 | 40.08 | 47.40 | 37.32 |
| L1 | 30.35 | 19.13 | 31.92 | 18.60 | 62.27 | 37.73 | 47.76 | 40.83 | 44.30 | 23.45 |
| L2 | 31.10 | 22.01 | 30.82 | 16.07 | 61.91 | 38.09 | 46.53 | 52.17 | 49.35 | 15.07 |
| Avg | 28.95 | 19.78 | 31.83 | 19.44 | 60.78 | 39.22 | 45.83 | 41.80 | 43.82 | 27.49 |

### 3.5 The effect of mutation and natural selection pressure on CUB of HPV16

ENC plot was used to measure the relative effects of mutational pressure and natural selection on CUB. In Figure 2, the curve represents the expected ENC determined by GC3 content and the points represent the actual ENC values of the eight ORFs. Almost all ENC values of HPV16 ORFs lie below the standard curve, suggesting that in addition to mutation pressure, other factors, such as natural selection, also influence the codon usage pattern of HPV16. The mean ENC values for the HPV16 ORFs was 41.27, with seven out of the eight ORFs had ENC larger than 35, indicating that the overall extent of CUB in HPV16 genomes was low. Interestingly, E4, E5 and E7 exhibited relatively lower ENC than expected, especially the E5 ORF (the mean ENC value was 24.95), implicating relatively high CUB. Although ENC is generally independent of gene length, these may still be influenced by the extremely short length of the three ORFs (E4, 95aa; E5, 78aa; E7, 98aa).

**Figure 2.**
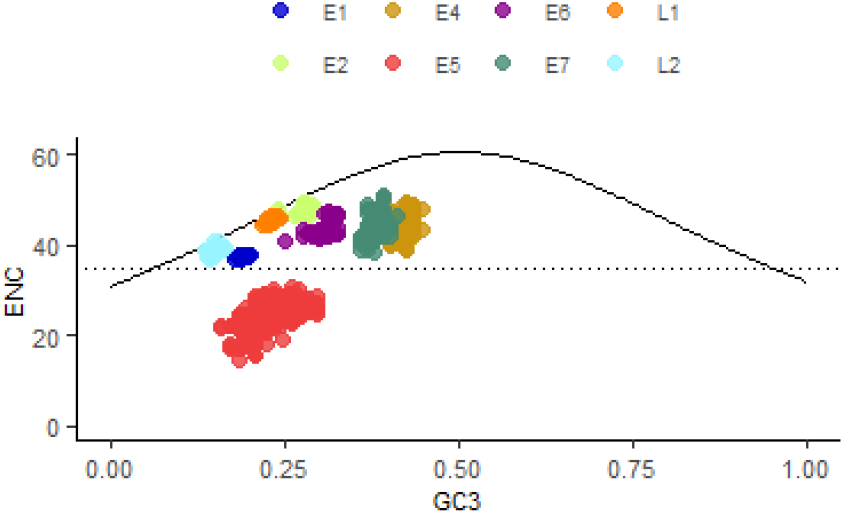
ENC plot of the eight ORFs of HPV16. The continuous curve plots the relationship between GC3 and ENC in the absence of selection. The horizontal dotted line represents the ENC value of 35. All points lie below the expected curve.

To further understand the influence of mutational or translational selection in HPV16 codon usage, regression analysis was conducted using GC12 (the mean GC content at the first and second codon positions) and GC3 (GC content of the third codon position) of each ORF (Figure 3). We observed a high correlation between GC12 and GC3 of E5 (R^2^=0.84), indicating mutation pressure on all the three codons. However, as previously mentioned, this result might be partly influenced by the short length of this gene. For the remaining seven ORFs, we didn’t observe high correlations (R^2^ <0.5). Most of the ORF (E1, E2, E4, E5, E6 and L1) were partly (15% to 30%) influenced by mutational pressure except for E7 and L2. L2 was found to be largely controlled by mutational pressure (81%). E7 was minimally (4%) governed by mutational pressure, indicating that the expression of this protein was largely affected by translational pressure. We have to point out that, because our data was skewed toward the A1 sublineage, the regression analysis might be biased.

**Figure 3.**
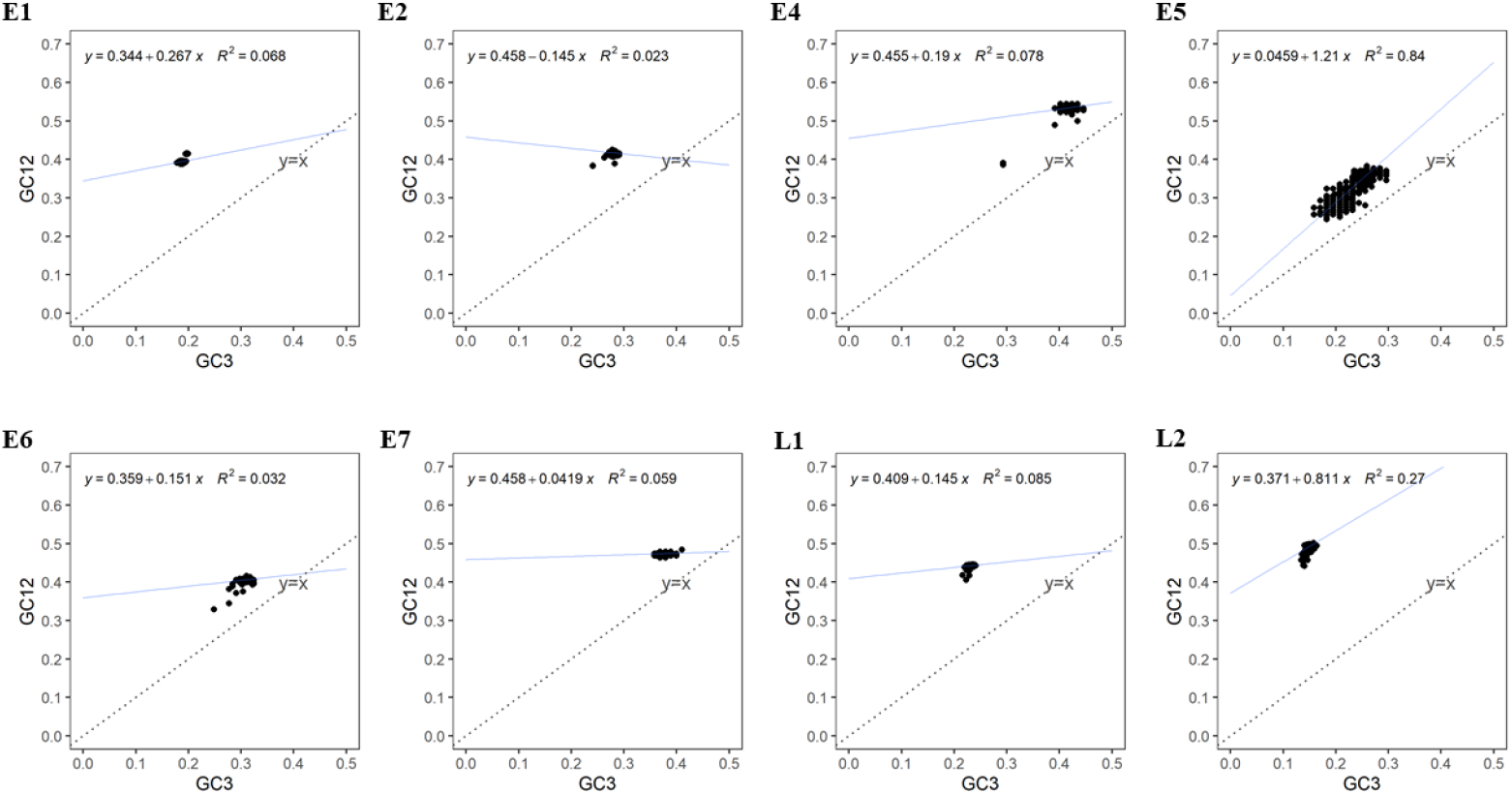
Neutrality plot analysis of GC12 and GC3 for HPV16 ORFs

### 3.6 Analysis of RSCU

To measure the usage variations of each codon, we calculated the RSCU values for HPV16 ORFs. The RSCU values varied in the eight ORFs (Figure 4). The RSCU of most codons ending in G/C was below 0.6, indicating that the usage frequency of these codons was relatively low. In contrast, RSCU values greater than 1.6 were mostly found in codons ending in A/T, indicating high usage preference. The top highly used codons including GCA for alanine, CCA for proline, ACA for threonine, TTA for leucine, AGA for arginine. TTA (leucine) was both highly used in L1 and L2 gene, AGA was the highly used codon in E6 gene, while E7 gene mostly preferred the codon of GTA (Supplementary Table S5). This finding was consistent with the high AT content in the nucleotide composition of the ORFs.

**Figure 4.**
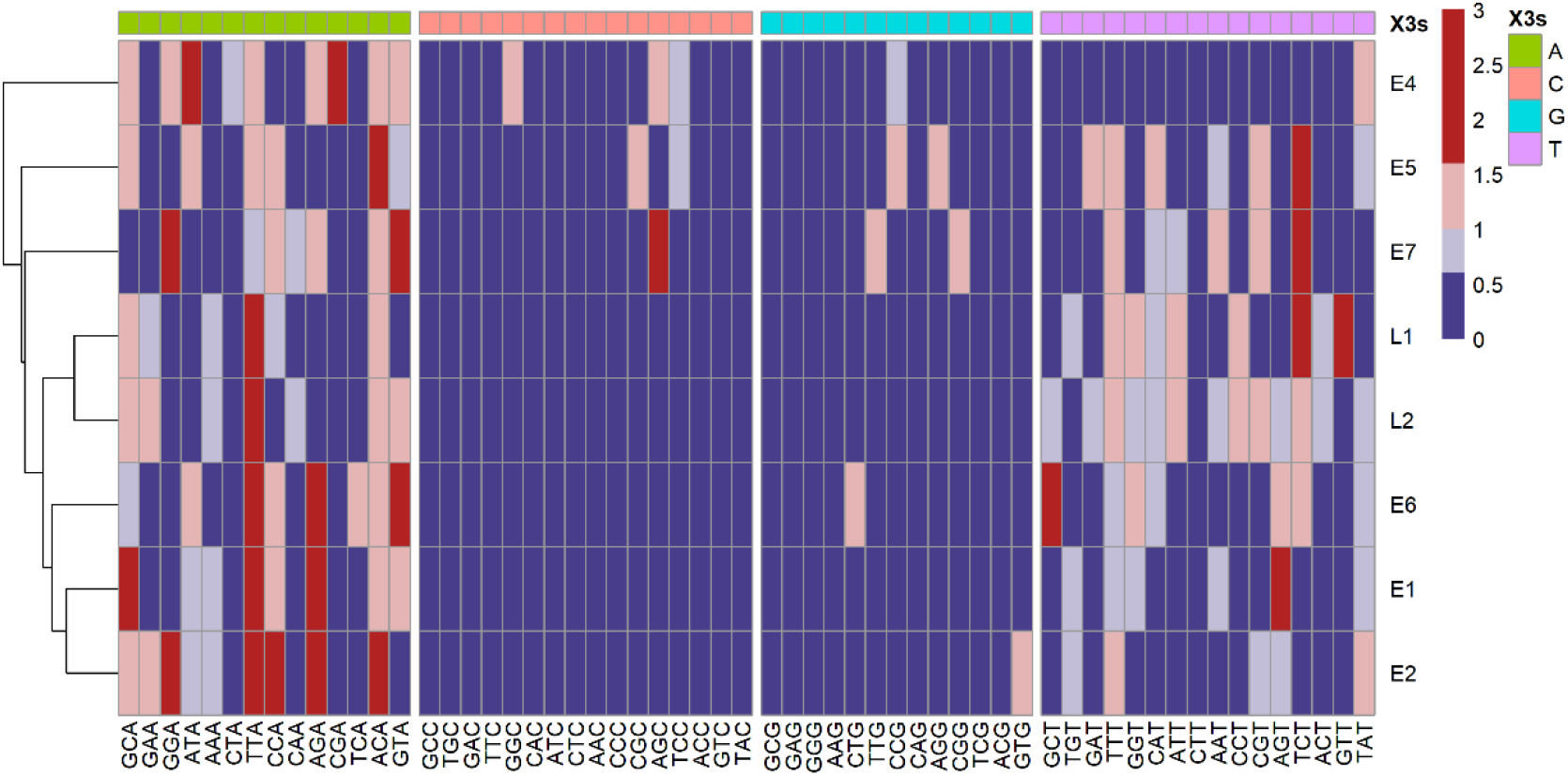
Relative synonymous codon usage (RSCU) analysis revealed over-representation of codons ending in A/T in HPV16 ORFs. Columns correspond to the 59 codons (three stop codons and those for Trp, Met were excluded). Rows correspond to the eight ORFs. Blue cells indicate under-represented codons (RSCU < 0.6) and red cells indicate over-represented codons (RSCU > 1.6). “X3s” indicated the 3^rd^ position nucleotide of synonymous codon

To understand the codon usage compatibility between virus and host, a correlation analysis between RCSU values of the eight HPV16 ORFs and those of humans was performed (Figure 5). The low R square values indicated that the codon usage preferences of the two species were only partially overlapped, with around 22-35 commonly preferred codons (i.e., normal and over usage) and 3-5 commonly preferred codons (Figure 5, bottom panel). These left 14-27 codons that were only preferred by human and 5-7 codons only preferred by HPV16. These results suggested that HPV16 was adapted in using the host translational machinery, but also avoided over competition with cellular protein production to reduce stimulation of the host immune response, which would help its persistence in human.

**Figure 5.**
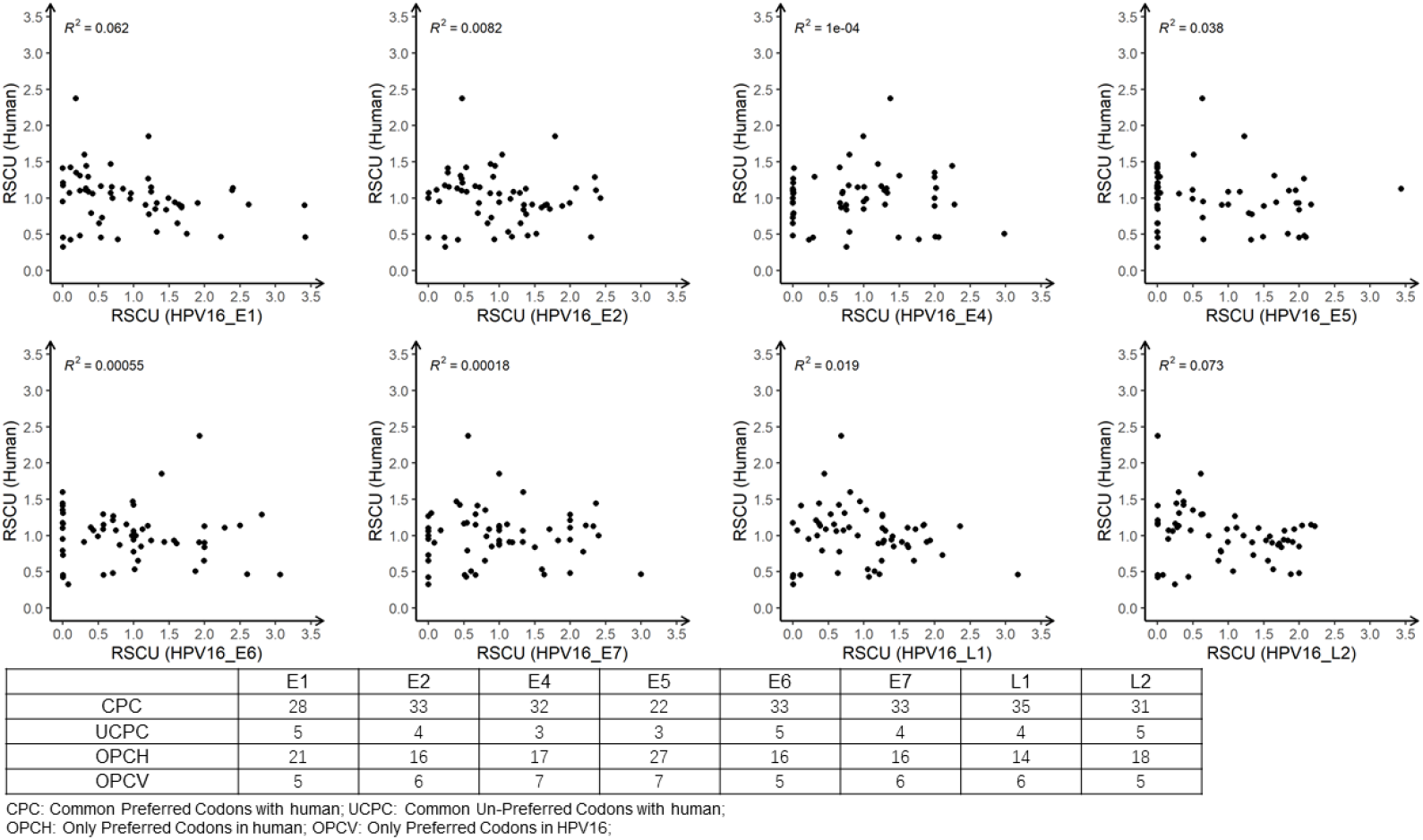
Pairwise correlation analysis of RSCU for 59 codons in eight HPV16 ORFs versus those of human. The R-squared values of linear regression analysis are shown. The embedded table denotes the number of common preferred (RSCU ≥ 0.6) codons and unpreferred (RSCU < 0.6) codons for the eight ORFs of HPV16 with human, and the number of preferred codons in human but unpreferred in HPV16 and preferred codons in HPV16 but unpreferred in human.

## 4. Discussion

Mutations in viral genes are important for variant identification and functional annotation. In our results, the most common mutations were T350G in the E6 gene and A647G in the E7 gene (Table 1). It was reported that these two mutations were related with the development of disease [48–50] and may be more common in China [51]. The HPV16 E6 T350G (L83V) variant has been shown to be prevalent in patients with high-grade cervical lesions and was strongly associated with cervical cancer progression [52, 53]. While A647G on the HPV16 E7 gene has been seen in other reports and it is thought that the mutation may be associated with persistent infection [49, 50]. Our mutation analysis showed that T350G mutation was detected in all viruses of lineage D and some strains in A1, A2 sublineages, while E7 A647G was observed in almost all A4 and C1 sublineages. This indicated that these mutations were not lineage or sublineage-specific. HPV16 E6 D25E was associated with an elevated risk for development of invasive cervical cancer [54]. Kahla et al. [55] reported that the mutation T310K in HPV16 E2 reduces the E2 DNA binding affinity and reverses its transcriptional regulatory activity on the early promoter of the virus. However, these two mutations were not identified in this study, possibly due to their scattered distribution across the different sublineages. We also found some lineage/sublineage-specific nucleotide variations. For example, A4967G and A5032T were only observed in D lineage and C3158G was a specific mutation in D4 sublineage. Lineage/sublineage-specific variants are highly correlated and represent fixed changes. These lineage/sublineage-specific mutation maybe helpful to determine the different HPV16 lineages/sublineages of infections.

Glycosylation modification of viral surface proteins is critical for viral infectivity and antigenicity, which has been documented for influenza viruses [26], Dengue viruses [56], HIV viruses [24], etc., and is a factor to be considered in the design of highly immunogenic vaccine. Among the four HPV16 lineages, lineage D contained the largest number of different glycosylation sites in L1 and L2 proteins from lineage A (Figure 1). Godi et al showed that comparing with HPV16 lineage A, lineage B, C, and D exhibited slightly (<2-fold) reduced sensitivity to nonavalent vaccine sera [57]. The unique glycosylation sites existed on the L1 proteins of lineages B, C and D, especially D, might be one of the determinants for this difference. Importantly, Zhou et al. reported that glycosylated L1 remained in the endoplasmic reticulum and was not transported for viral particle assembly, suggesting that glycosylated L1 might not be an important component of the papillomavirus virion [58]. Additional studies are needed to demonstrate the function of glycosylation sites of HPV16 L1 and L2 proteins and the impact of glycosylation on the design of HPV vaccines.

Our nucleotide composition analysis showed that the A+T content of HPV16 was higher than the G+C content in most HPV16 ORFs. Zhao et al. [32] analyzed 79 HPV types and showed that the E4 gene was GC-rich while the other open reading frames were AT-rich, this result was in accordance with our findings. It has been shown that GC3 was associated with the CUB of the organism [59–61], GC-rich codons were more likely to end in GC, and vice versa. We found that the GC3 content varied greatly between different ORFs of HPV16, ranging from 15.07% to 41.85%, which was closely related to codon usage preference. Consistently, we found that the relative synonymous codon usage was higher for codons ending in AT. In our analysis, the ENC values of the HPV16 genes were above 35 except that of E5 gene, indicating a lower codon preference and possibly low gene expression level [59, 62]. The statement that ENC calculation was generally independent of gene length was true for genes with over 100 codons but may not be applicable for short genes [63]. Therefore, the ENC results for the three ORFs (E4, E5 and E7) with less than 100 codons should not be over-interpreted. The CUB of organisms is largely influenced by natural selection and mutational pressure [60, 64, 65]. Our ENC and neutrality results indicated that the main factor affecting HPV16 codon usage might be translational selection, except for E5 and E7 genes. We also found that the codon usage of HPV16 did not fully overlap with that of humans, which might help the virus better evade host immunity to facilitate persistent infection in human.

Using a large amount of HPV16 genomes (1,597), we have comprehensively investigated the mutation profiles across the HPV16 genome, potential glycosylation site distribution in surface proteins and the codon usage patterns of all the eight ORFs of HPV16. These findings might provide important implications for variant identification, novel vaccine development and give hints on the viral-host interaction mechanism supporting the chronic viral infection in humans. Currently the available HPV16 genomes were mainly from lineage A, especially sublineage A1. Therefore, our neutrality plot might be greatly affected by the abundant similar sequences. Increased genomic surveillance around the world may help reveal the complete sublineage diversity of HPV16 and improve the genomic research on the viruses.

## Supporting information

all supplemental files

## Supplementary Materials

The following are available online at http://www.mdpi.com/xxx/s1,

Table S1:The detailed information of HPV16 genomes downloaded from public database.

Table S2:Summary of the lineage/sublineage distribution of HPV16 genomes.

Table S3:All mutations observed in HPV16 ORFs.

Table S4:Potential glycosylation sites in L1 proteins of HPV16 sublineages.

Table S5: Potential glycosylation sites in L2 proteins of HPV16 sublineages.

Table S6:The RSCU values of 59 synonymous codons in eight HPV16 ORFs.

Figure S1: Phylogeny of HPV16 complete genomes. Maximum likelihood phylogeny was constructed with IQ-TREE using TVM+F+I+G4 nucleotide substitution model. Bootstrap values over 70 were labelled in purple. The tree scale was displayed at the bottom. The pairwise nucleotide sequence differences were calculated for each isolate and are shown on the right panel. The references genomes were labelled by black solid circles. Different colors indicated different lineages/sublineages.

Figure S2: Mutation distribution across the HPV16 genome.

## Author Contributions

Z.O. designed and supervised the investigation; W.L. conducted literature review, performed data analysis and visualization; W.L. and Z.O. prepared the manuscript. J.L. and H.D. provided critical advice for the manuscript. All authors have read and agreed to the published version of the manuscript.

## Acknowledgments

We thank all members of the Infection Omics Research Center for their instructive academic advice. Miss Wei Liu would like to express gratitude to her beloved families, Miss Qing Nie and Mr. Zhaohui Shen. Dr. Zhihua Ou would like to thank the warm support from Miss Feiyun Ou and Mr. Geer Xi.

