## Supplementary figures and images for "Mutation Profiles, Glycosylation Site Distribution and Codon Usage Bias of HPV16"

### Figure S1_hpv16_gc_selected_final.fasta.treefile_dis.pdf

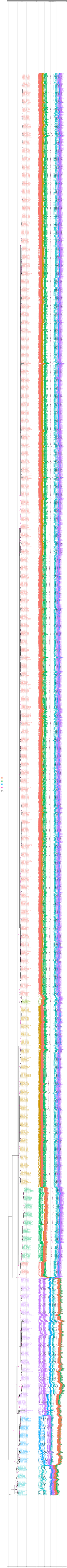

### Figure S2_hpv16_mutation.tif

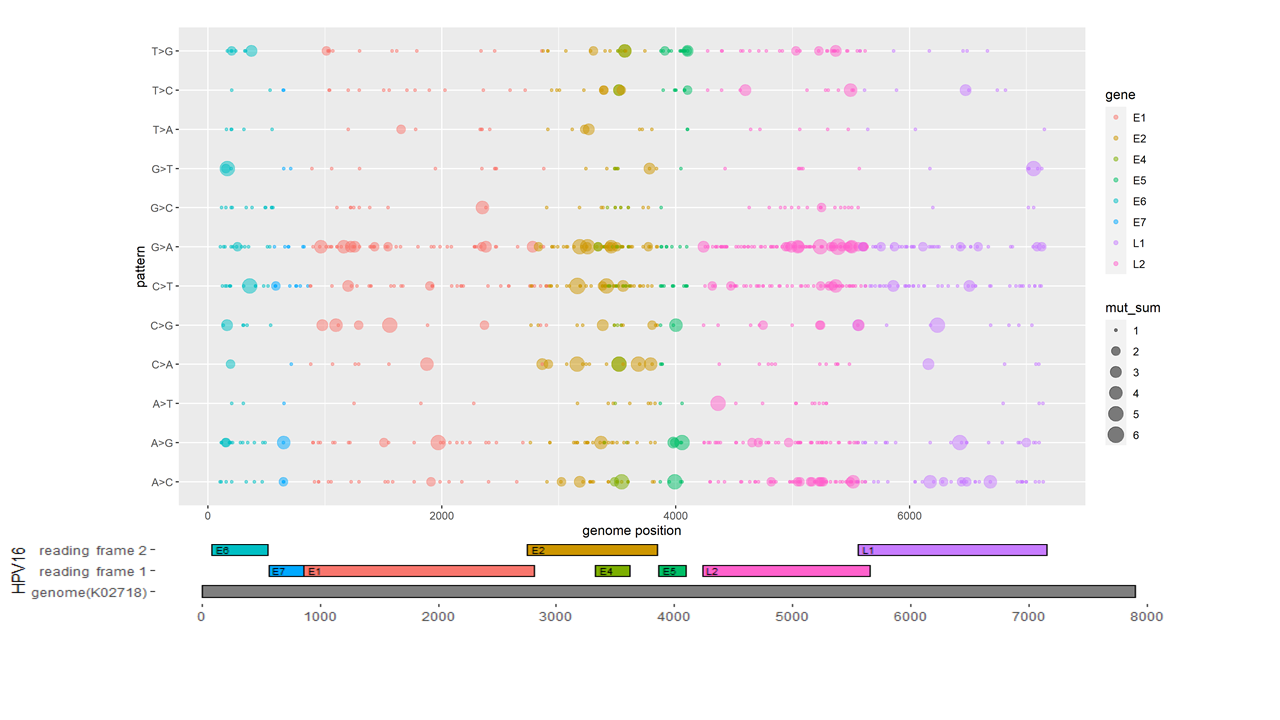
